# A Generalized Robust Allele-based Genetic Association Test

**DOI:** 10.1101/2020.03.12.989004

**Authors:** Lin Zhang, Lei Sun

## Abstract

The allele-based association test, comparing allele frequency difference between case and control groups, is locally most powerful. However, application of the classical allelic test is limited in practice, because the method is sensitive to the Hardy–Weinberg equilibrium (HWE) assumption, not applicable to continuous traits, and not easy to account for covariate effect or sample correlation. To develop a generalized robust allelic test, we propose a new allele-based regression model with individual allele as the response variable. We show that the score test statistic derived from this robust and unifying regression framework contains a correction factor that explicitly adjusts for potential departure from HWE, and encompasses the classical allelic test as a special case. When the trait of interest is continuous, the corresponding allelic test evaluates a weighted difference between individual-level allele frequency estimate and sample estimate where the weight is proportional to an individual’s trait value, and the test remains valid under *Y* - dependent sampling. Finally, the proposed allele-based method can analyze multiple (continuous or binary) phenotypes simultaneously and multi-allelic genetic markers, while accounting for covariate effect, sample correlation and population heterogeneity. To support our analytical findings, we provide empirical evidence from both simulation and application studies.

## 1 Introduction

An association study aims to identify genetic markers that influence a heritable trait or phenotype of interest, while accounting for environmental effect. To formulate the problem more precisely, let single nucleotide polymorphisms (SNPs) be the genetic markers available for analysis. For each bi-allelic SNP, let *a* and *A* be the two possible alleles, and as in convention, let *A* denote the minor allele with population frequency *p* ≤ 0.5. The SNP genotype consists of a paired (but unordered) alleles, *aa*, *Aa* or *AA*. For a case-control association study of a binary trait (Table 1), intuitively one can compare the estimates of *p* between the case and control groups. Indeed, the resulting allelic test is locally most powerful (Sasieni, 1997). However, the validity of the test hinges on the assumption of Hardy–Weinberg equilibrium (HWE), which assumes that genotype frequencies depend only on allele frequencies. That is, *p*_*aa*_ = (1 − *p*)^2^, *p*_*Aa*_ = 2*p*(1 − *p*) and *p*_*AA*_ = *p*^2^.

**Table 1:** Notations for genotype and allele counts for a case-control study. The HLA-DQ3 example is from Sasieni (1997), studying women with cervical intraepithelial neoplasia 3. The sample estimates of allele frequency of allele A in the case, control and combined groups are denoted, respectively, as 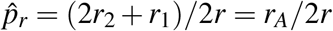, 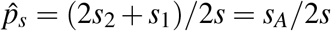 and 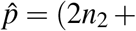 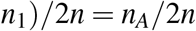

|  | Genotype Counts |  |  |  | Allele Counts |  |  |
| --- | --- | --- | --- | --- | --- | --- | --- |
| | $aa$ | $Aa$ | $AA$ | Total | $a$ | $A$ | Total |
| Case | $r_0$ | $r_1$ | $r_2$ | $r$ | $2r_0 + r_1$ | $r_1 + 2r_2$ | $2r$ |
| Control | $s_0$ | $s_1$ | $s_2$ | $s$ | $2s_0 + s_1$ | $s_1 + 2s_2$ | $2s$ |
| Total | $n_{aa}$ | $n_{Aa}$ | $n_{AA}$ | | $n_a$ | $n_A$ | |
| | $n_0$ | $n_1$ | $n_2$ | $n$ | $2n_0 + n_1$ | $n_1 + 2n_2$ | $2n$ |
| The HLA-DQ3 example from Sasieni (1997) |  |  |  |  |  |  |  |
| Case | 40 | 45 | 28 | 113 | 125 | 101 | 226 |
| Control | 273 | 100 | 43 | 416 | 646 | 186 | 832 |
| Total | 313 | 145 | 71 | 529 | 771 | 287 | 1058 |

To evaluate the HWE assumption in an independent sample, one typically applies the Pearson goodness-of-fit *χ*^2^ test, and under the *H*_0_ of HWE,

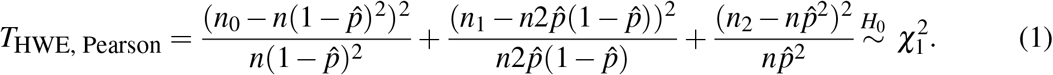

Using the HLA-DQ3 data in Table 1 as an illustration, application of *T*_HWE, Pearson_ yields a test statistic of 49.7623 and a p-value of 1.74 × 10^−12^, suggesting departure from HWE.

Prior to conducting a case-control genome-wide association study (GWAS), testing HWE in the control sample and excluding SNPs in Hardy–Weinberg disequilibrium (HWD) is a routine part of data quality control; severe HWD is an indication of genotyping error. However, the need for a robust allelic association test remains. First, the p-value threshold used for HWE-based screening in practice is subjective and depends on sample size, often ranging between 10^−8^ and 10^−12^. For example, Bycroft et al. (2018) recommended 10^−12^ for each batch of ~4,000 samples. With this stringent criterion, it is possible that a SNP that passes the quality control step is truly in HWD at the population level, but not attributed to genotyping error. The corresponding allelic test is then biased, and we provide an application example in Section 5.2. Second, developing a robust and generalized allelic test is analytically important, so that this locally most powerful test (Sasieni, 1997) can be applied to, for example, a continuous trait or a sample of related individuals.

To robustify the classical allelic test against HWD, early work focused on improving variance estimation (Schaid and Jacobsen, 1999), but application is still limited to case-control studies and in simple settings (e.g. independent samples). Thus, most genetic association studies rely on the *Y* -on-*G* regression approach, where *G* is coded additively as 0, 1 and 2 for *aa*, *Aa* and *AA*. The resulting genotype-based additive test is known to be robust to HWD, but the exact HWD-correction mechanism is not clear. Further, data collection in practice typically first samples individuals based on *Y*, which can be random or *Y* - dependent sampling (Derkach et al., 2015), and then collects *G* for the sampled individuals. Thus, it can be argued that *G*-on-*Y* is a more fitting statistical framework.

This ‘reverse’ regression approach can also readily analyze multiple phenotypes simultaneously, which was the motivation for MultiPhen (O’Reilly et al., 2012). To deal with the three genotype groups, O’Reilly et al. (2012) used an ordinal logistic regression and stated that the proposed likelihood ratio test does not assume HWE. However, the statistical insight is lacking and analyzing pedigree data remains a challenge. Zhang et al. (2014) provided more background on using ‘reverse’ regression to simultaneously analyze multiple traits, and also proposed and compared a series of genotype-based GEE association tests.

This work generalizes the locally most powerful allelic association test to more complex settings by developing a novel *allele-based* ‘reverse’ regression model. Section 2 revisits the classical allelic association test and provides insight on the need for a more flexible formulation of the test. Section 3 develops the new allele-based ‘reverse’ regression model by first appropriately partitioning the two alleles of a genotype and then specifying the individual allele as the response variable. In addition to the association parameter, the proposed regression includes a parameter that models the dependency between the two alleles of a genotype, explicitly accounting for potential departure from HWE. Section 4 considers more complex settings including related individuals from pedigree data, genetic markers with more than two alleles, and multiple phenotypes or populations. Given the theoretical results presented, simulation experiments in Section 5 are relatively brief with additional empirical evidence from two applications. Section 6 concludes with remarks and discussion.

## 2 The classical allelic test revisited

For a given SNP and a binary phenotype of interest, let *p*_*r*_ and *p*_*s*_ denote the allele frequencies, respectively, for the case and control populations (Table 1). The classical allele-based association test, testing *H*_0_: *p*_*r*_ = *p*_*s*_, is based on

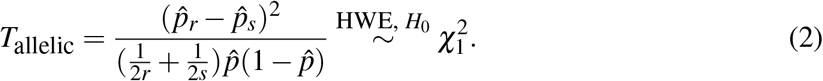

The validity of *T*_allelic_, however, requires the Hardy–Weinberg equilibrium assumption, because only under HWE, 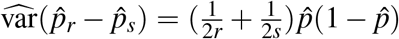. Using the HLA-DQ3 data in Table 1 as an example, because the HWE test is significant (Section 1), a direct application of the classical allelic association test in this case is not appropriate.

Indeed, Sasieni (1997) has pointed out that *T*_allelic_ is valid and locally most powerful if and only if the HWE assumption holds and the genetic effect is additive. In the presence of Hardy– Weinberg disequilibrium, *T*_allelic_ is known to be anti-conservative. But, we emphasize that this is true only if there is an excess of homozygotes *AA*, i.e. *δ* = *p*_*AA*_ − *p*^2^ *>* 0, where *δ* is a measure of HWD (Weir, 1996). If *δ <* 0, *T*_allelic_ is conservative as shown in Figure 1.

**Figure 1:**
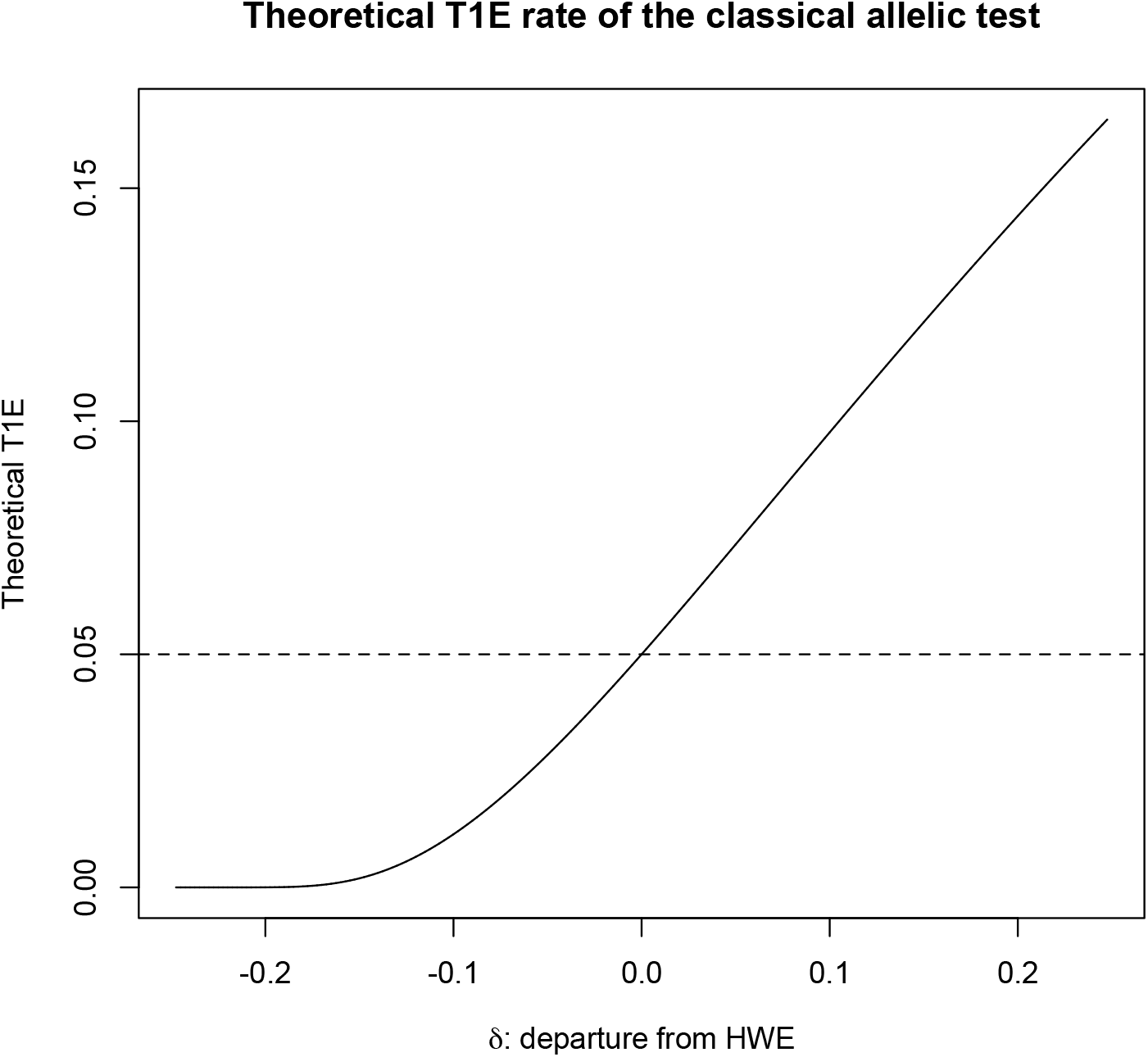
The theoretical type 1 error rate of the classical allelic test, *T*_allelic_. The nominal level is *α* = 0.05, departure from HWE is measured by *δ* = *p*_*AA*_ − *p*^2^, where *p* is the frequency of the minor allele *A* and −*p*^2^ ≤ *δ* ≤ *p*(1 − *p*). When *p* = 0.5, −0.25 ≤ *δ* ≤ 0.25.

To robustify *T*_allelic_ against HWD, Schaid and Jacobsen (1999) used a multinomial distribution for the genotype counts to improve variance estimate. The resulting test is

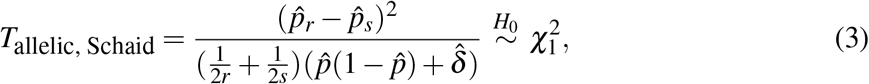

where 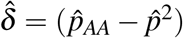 is the sample estimate of the HWD measure. Section 3 provides analytical insight about how *T*_HWE, Pearson_ in (1) is related to *δ*. Here, the effect of *δ* ≠ 0 on the accuracy of *T*_allelic_ is clear. In the HLA-DQ3 example, 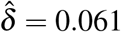 and is highly significant, so the classical allelic test will be too optimistic, {*T*_allelic_ = 44.8470} *>* {*T*_allelic, Schaid_ = 34.3207}.

*T*_allelic, Schaid_ is effective in making *T*_allelic_, the classical allele-based association test, robust to HWD, but its direct application is limited to the simplest setting of case-control studies using independent observations without covariates. It is also not clear how to adapt this comparing-two-proportions analytical framework to, for example, analyzing a continuous trait. Thus, a new formulation of allele-based association test is needed.

## 3 A Generalized Robust Allele-based (RA) Association Test

### 3.1 Decoupling the two alleles in a genotype

Consider a bi-allelic SNP with genotype *G* ∈ {*aa, Aa, AA*}, and for the moment assume that there are *n* independent observations, *G*_*i*_, i = 1*, ..., n*. We partition each *G*_*i*_ as follows,

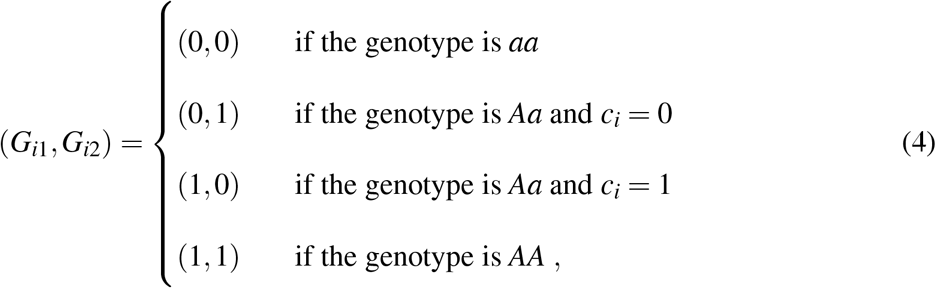

where 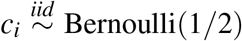 if *G*_*i*_ = *Aa*. The partition of a genotype is straightforward for homozygotes *aa* or *AA*, but not so for a heterozygote *Aa*. Previous work attempted to split the *n*_*Aa*_ observations to (0, 1) and (1, 0) equally (Schaid et al., 2012; Bourgain et al., 2003), which reduces the variation inherent in a randomly selected allele (Web Appendix A). The use of a fair coin in our proposed approach ensures that ∑_*i*_ *G*_*i*1_ ~ Binomial(*n, p*_*AA*_ + *p*_*Aa*_/2) and similarly for ∑_*i*_ *G*_*i*2_, which is critical to valid HWE test as well as association inference.

### 3.2 Reformulating the test of HWE as an allele-based regression

A critical component of developing a robust allelic association test is the modeling of the Hardy–Weinberg equilibrium assumption. HWE assumes that the two alleles in a genotype are independent of each other, so a natural approach is to use a logistic regression, 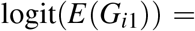 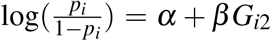. Indeed, we show in Web Appendix B that the corresponding score test of testing *H*_0_ : *β* = 0 approximates *T*_HWE, Pearson_; regressing *G*_*i*2_ on *G*_*i*1_ leads to the same conclusion. However, the differential treatment of *G*_*i*1_ and *G*_*i*2_ is not ideal. Further, the sample size of this model is *n* whereas there are 2*n* alleles from *n* genotypes. Thus, we consider an alternative regression model that ‘doubles’ the sample size.

The proposed robust allele-based (RA) regression model for HWE testing is

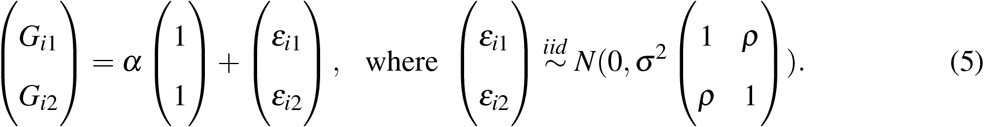

This RA model has two important features. First, instead of using the location parameter *β* to represent the dependence between two alleles, we re-parameterize it as the correlation parameter *ρ* in the covariance matrix to capture HWD. This model reformulation is crucial to methodology development in Section 3.3 where the regression coefficient is reserved for association testing. Second, instead of a logistic regression, we use a Gaussian model even though the outcome is binary. This is because the primary goal here is hypothesis testing not parameter estimation or interpretation. For testing the *location* parameters in regression, Chen (1983) has shown that, under some regularity conditions, the score test statistics for regression models from the exponential family have identical form. In our setting, the Gaussian model enables explicit modeling of HWD through the *correlation* parameter *ρ*, and the resulting score test encompasses *T*_HWE_, Pearson as a special case which we show below.

Based on the Gaussian model of (5), the score test statistic of testing *H*_0_ : *ρ* = 0 is

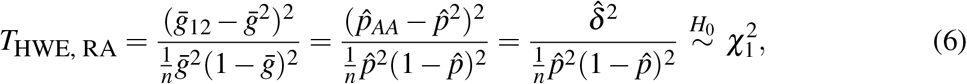

where 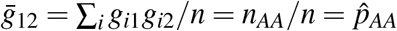, 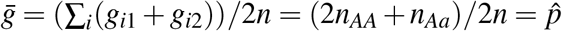, and 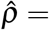 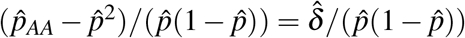, which is a scaled estimate of HWD, *δ*.

We first note that the newly developed HWE test statistic is, attractively, proportional to 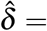 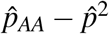. Interestingly, after some algebraic manipulations, we show in Web Appendix C that *T*_HWE, RA_ in (6) is *identical* to *T*_HWE, Pearson_ in (1). This equivalence, however, is under the simplest scenario of an independent sample. For more complex data, several authors have proposed different HWE testing strategies, e.g. Troendle and Yu (1994) developed a method that tests HWE across strata, while Bourgain et al. (2004) proposed a quasi-likelihood method that tests HWE in related individuals. Section 4 shows how the proposed regression (5) can be extended to derive a generalized HWE test. For the moment, we still consider an independent sample but turn our attention to association analysis.

### 3.3 The generalized robust allele-based (RA) association test via regression

Let *G*_*i*1_ and *G*_*i*2_ be the two allele-based random variables, as constructed in (4) for a bi-allelic SNP in an independent sample of size *n*, *i* = 1*, ..., n*. We now also consider *Y*, a (categorical or continuous) phenotype of interest, and *Z*, one single covariate for notation simplicity but without loss of generality. The proposed RA regression for association analysis is,

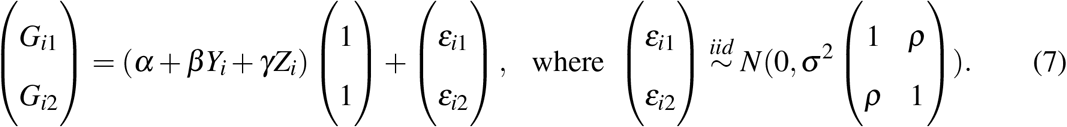

The corresponding score test, testing *H*_0_ : *β* = 0, is

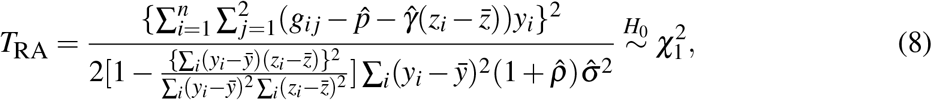

where 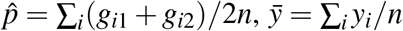, 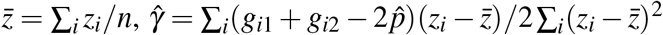, 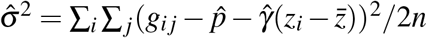, and 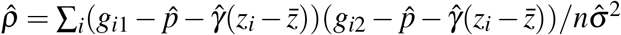. We provide the general form of *T*_RA_ with multiple *Y* s and multiple *Z*s in Web Appendix D.

The proposed *T*_RA_ unifies previous methods. For example, if *Y* is binary and there are no covariates, *T*_RA_ in (8) is simplified to

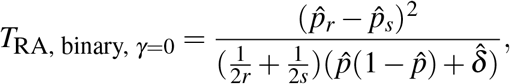

which is *T*_allelic, Schaid_ in (3). If we further assume HWE, *T*_RA_ is reduced to

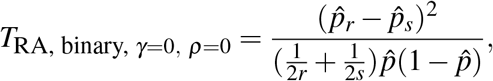

which is the classical allelic test, *T*_allelic_ in (2).

The proposed RA test also generalizes the allelic association test. To provide analytical insight, consider a *continuous* trait without covariate effect. In that case,

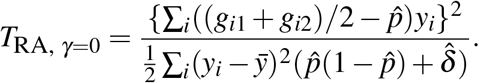

Thus, for a continuous trait, the generalized allelic association test evaluates a weighted difference between individual-level allele frequency estimate, (*g*_*i*1_ + *g*_*i*2_)/2, and the whole sample estimate, 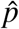, where the weight is an individual’s trait value, *y*_*i*_. In hindsight, this result may not be surprising. However, the advantages of developing the proposed RA regression model become evident when we analyze more complex data such as data with population heterogeneity and related individuals, which we investigate in the next section.

## 4 Complex data

### 4.1 Multiple populations, alleles, or phenotypes

The proposed RA regression model of (7) can naturally adjust for population effects by including population indicators, or top principal components, as part of the covariates. Here we emphasize that the potential population effects could include both difference in allele frequency and difference in HWD between populations. The RA framework models allele frequency heterogeneity through *γ* and *σ*^2^, while accounting for HWD through *ρ*.

Without loss of generality, it is instructive to consider the simple case of a case-control study with two populations but without additional covariates. Denote *Z*_*i*_ = 0 for population I and *Z*_*i*_ = 1 for population II. The corresponding RA regression model is

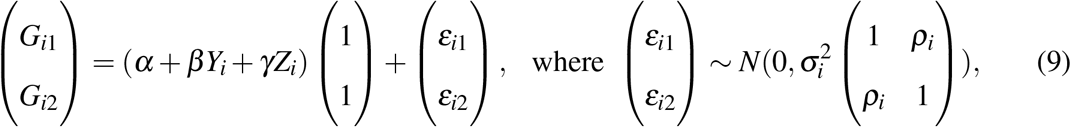

*ρ*_*i*_ = *ρ*^*I*^ and 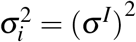 if *Z*_*i*_ = 0; *ρ*_*i*_ = *ρ*^*II*^ and 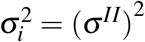 if *Z*_*i*_ = 1. Using superscripts ^*I*^ and ^*II*^ for all the other notations introduced so far, the generalized RA test of *H*_0_ : *β* = 0 while accounting for population heterogeneity has the following expression,

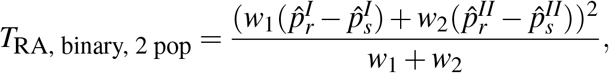

where 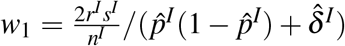, and 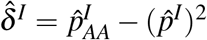 captures population-specific HWD in population I; the analytical expressions are the same for *w*_2_ and 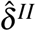.

If evaluating HWE across multiple populations is the primary objective, we can test *H*_0_ : *ρ*^*I*^ = *ρ*^*II*^ = 0 and show that *T*_HWE, RA, 2 pop_ = *T*_HWE, RA, pop I_ + *T*_HWE, RA, pop II_ ~ 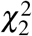, where the expressions for *T*_HWE, RA, pop I_ and *T*_HWE, RA, pop II_ are given in (6). We note again the unifying feature of the proposed RA framework. For example, the test of Troendle and Yu (1994) developed specifically for testing HWE across strata has identical form as *T*_HWE, RA, 2 pop_.

The RA model of (7) can be also extended to derive a generalized allelic association test for multi-allelic markers with *K >* 2 alleles, with adjustments for covariate effect and Hardy–Weinberg disequilibrium. In that case, we need to introduce two indicator vectors, *g*_*i*1_ and *g*_*i*2_, each of length *K* − 1, to partition the genotype of individual *i*; we leave the technical details to Web Appendix E and Web Table 1. Given *g*_*i*1_ and *g*_*i*2_, the implementation of the RA regression model of (7) is straightforward. The RA model can also be used to derive regression-based HWE test of a multi-allelic marker as shown in Web Appendix F.

To analyze multiple *J* phenotypes simultaneously, we can simply include multiple *Y*_*j*_1 vectors in the RA model of (7), each representing one phenotype, and then test *H*_0_ : *β*_*j*_ = 0, ∀ *j* ∈ {1, 2*, ..., J*}. The corresponding score test statistic is 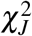 distributed under the null of no association. Here we re-iterate that the proposed ‘reverse’ regression is *allele-based*, conceptually distinct from the *genotype-based* MultiPhen (O’Reilly et al., 2012) which uses an ordinal logistic regression for the three genotype groups.

### 4.2 Related individuals

We now consider a sample of *n* correlated individuals with known or accurately estimated pedigree structure (Dimitromanolakis et al., 2019). For notation simplicity but without loss of generality, we present results for analyzing a bi-allelic SNP and one phenotype in a homogeneous population. Let *g* be a 2*n* × 1 vector of allele indicators for the *n* genotypes available, where 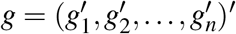 and *g*_*i*_ = (*G*_*i*1_, *G*_*i*2_)′ for *i* ∈ {1*,..., n*}, following the allele-partition method in (4), and let 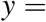 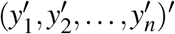, 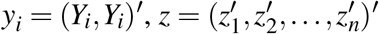, and 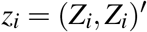. The generalized RA regression model for a dependent sample is,

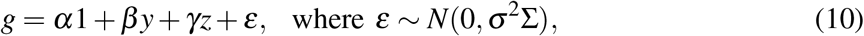

1 is a 2*n* × 1 vector of 1s, and Σ is a 2*n* × 2*n* matrix that captures the genetic correlation between individuals as well as departure from Hardy–Weinberg equilibrium in founders. Founders are individuals who have no ancestors or siblings included in the sample, and their offspring genotypes are in HWE assuming random mating (Web Appendix G).

The specification of Σ is non-trivial, where for any two individuals *i* and *j*, 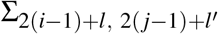 not only measures the genetic correlation between individual *i*’s *l*th allele and individual *j*’s *l*′th allele, *l* and *l*′ ∈ {1, 2}, but also accounts for potential HWD. We note that if *i* = *j* and *l* = *l*′, 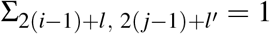. If *i* = *j* and *l* ≠ *l*′, 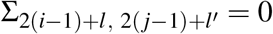 for a non-founder and = *ρ* for a founder, where *ρ* models HWD. Finally, if *i* ≠ *j*, 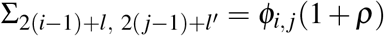, where *ϕ*_*i, j*_ is the kinship coefficient between the two individuals (Web Appendix H).

As an illustration, consider a sample of *f* independent sib-pairs. With a slight abuse of notations, let {*G*_*j*11_, *G*_*j*12_, *G*_*j*21_, *G*_*j*22_} denote the the four alleles of the *j*th sib-pair, *j* = 1*, ..., f*, where {*G*_*j*11_, *G*_*j*12_} are for sibling 1 and {*G*_*j*21_, *G*_*j*22_} are for sibling 2. In this case, Σ is a block diagonal matrix with

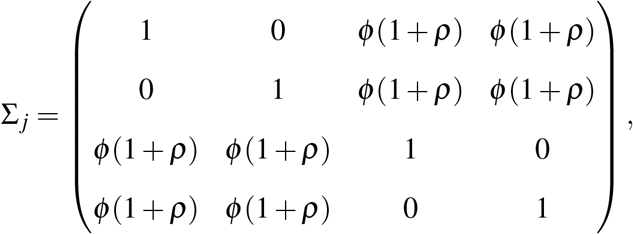

where *ϕ* = 0.25 is the kinship coefficient for a sib-pair.

For further illustration, assume all sib-pairs are concordant in phenotype (i.e. *r* pairs of cases and *s* pairs of controls) with no covariates, the score statistic of testing *H*_0_ : *β* = 0 is

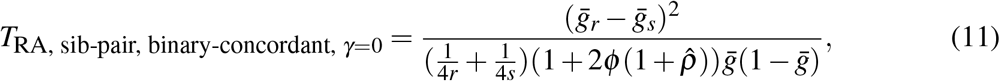

where 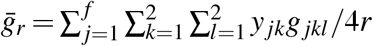, 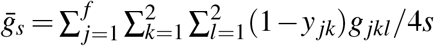, 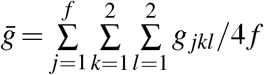 and 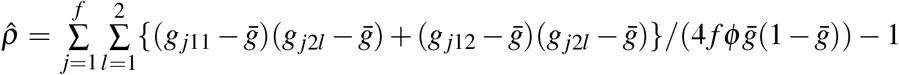. It is compelling that the expression of (11) is similar to that of the classical allelic test in (2). However, the denominator in (11) explicitly adjusts for the inherent genetic correlation *between* (not within unless inbreeding) the sibling alleles through *ϕ*, as well as any potential HWD through 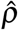.

## 5 Empirical evidence

### 5.1 Simulation studies

To numerically demonstrate the robustness of *T*_RA_ to Hardy–Weinberg disequilibrium as compared with *T*_allelic_, we simulated a case-control study with an independent sample of 1,000 cases and 1,000 controls. The minor allele frequency (MAF) was *p* = 0.2 or 0.5 for the minor allele *A*. The amount of HWD as measured by *δ* = *p*_*AA*_ − *p*^2^ ranged from the minimum of −*p*^2^ to the maximum of *p*(1 − *p*). Then *p*_*AA*_ = *p*^2^ + *δ* and *p*_*Aa*_ = 2*p*(1 − *p*) − 2*δ*, and (*n*_*aa*_, *n*_*Aa*_, *n*_*AA*_) ~ Multinomial{*n,* (1 − *p*_*Aa*_ − *p*_*AA*_, *p*_*Aa*_, *p*_*AA*_)}. For power evaluation at *α* = 0.05, we assumed an additive model with disease penetrances *f*_0_ = *P*(*Y* = 1|*G* = *aa*) = 0.08, *f*_1_ = *P*(*Y* = 1|*G* = *Aa*) = 0.10, and *f*_2_ = *P*(*Y* = 1|*G* = *AA*) = 0.12.

The empirical type 1 error results in Figure 2(a) and 2(b) confirm the theoretical results in Figure 1: the proposed *T*_RA_ is accurate across the whole range of HWD values, while *T*_allelic_ is not robust against HWD. Further, the empirical power results in Figures 2(c) and 2(d) highlight the fact that the classical allelic test could have reduced power when the number of homozygotes *AA* is fewer than what is expected under the HWE assumption (i.e. when *δ <* 0), which is not well acknowledged in the existing literature.

**Figure 2:**
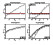
Empirical type 1 error rate and power of the classical allelic association test and the proposed robust allelic (RA) test. The nominal level is *α* = 0.05, evaluated based on 10^4^ simulation replicates. Note that when *δ >* 0, the classical allelic test has inflated type 1 error rate as shown in (a) and (b), so the corresponding power in (c) and (d) is not meaningful and shown in a lighter shade. Also note that the HWD measure *δ* is bounded by the minor allele frequency *p*, *p*^2^ *δ p*(1 − *p*). Results of type 1 error control when analyzing rare variants with *p* = 0.01 or 0.05 are shown in Web Figure 2, and when using the GWAS significance level of *α* = 5 10^−8^ (evaluated based on 10^10^ simulation replicates) are shown in Web Table 2.

We also assessed the accuracy of *T*_RA_ and *T*_allelic_ when analyzing SNPs with low MAF (*p* = 0.01 and 0.05), and results in Web Figure 2 are similar to that in Figure 2. In addition, we evaluated the performance of the tests using the GWAS significance level of *α* = 5 × 10^−8^, with 10^10^ simulation replicates for each parameter setting considered (Web Table 2); results for MAF *>* 0.1 are not shown, because the performance of *T*_RA_ is better for larger MAF as expected. Overall, the proposed RA test is accurate or slightly conservative for the whole range of HWD. In contrast, the classical allelic test has grossly inflated type 1 error rate up to 1.35 × 10^−5^ for *α* = 5 × 10^−8^ (Web Table 2).

**Table 2:** Results of application 2 - Previously reported SNPs. The p-values in this table differ from those in Sun et al. (2012) and Corvol et al. (2015), because the analyses here only included the Canadian samples and individuals with both phenotypes measured. Other details see legend to Figure 4.

| Top CF lung function associated SNPs from Corvol et al. (2015) |  |  |  |  |
| --- | --- | --- | --- | --- |
| Chr | SNP | Lung function only | | Lung function & MI simultaneously<br>$-\log_{10}P_{RA, \text{ lung \& MI}}$ |
| | | $-\log_{10}P_{LMM}$ | $-\log_{10}P_{RA}$ | |
| 3 | rs2246901 | 3.21 | **3.25** | 2.63 |
| 5 | rs3749615 | 3.25 | **3.27** | 3.57 |
| 6 | rs2395185 | 6.65 | **6.77** | 6.08 |
| 11 | rs10466455 | 5.84 | **5.86** | 4.84 |
| Top CF meconium ileus associated SNPs from Sun et al. (2012) |  |  |  |  |
| Chr | SNP | MI only | | Lung function & MI simultaneously<br>$-\log_{10}P_{RA, \text{ lung \& MI}}$ |
| | | $-\log_{10}P_{GLMM}$ | $-\log_{10}P_{RA}$ | |
| 1 | rs4077468 | 5.34 | **5.47** | 4.87 |
| 1 | rs7512462 | 4.54 | **4.82** | 4.56 |
| 1 | rs7419153 | 3.68 | **4.07** | 3.35 |
| 1 | rs12047830 | 3.10 | **3.20** | 2.63 |

### 5.2 Application 1 - revisit the study of Wittke-Thompson et al. (2005)

Studying Hardy–Weinberg disequilibrium not due to genotyping error, Wittke-Thompson et al. (2005) identified 60 bi-allelic SNPs from 41 case-control association studies. Focusing on association analysis of these 60 SNPs, we compared *T*_allelic_ with the proposed *T*_RA_ (Figure 3). As expected based on the simulation results in Figure 2, for SNPs with 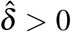, *T*_allelic_ can appear to be more powerful, due to increased type 1 error rate, than the proposed *T*RA. For example, for the most significant SNP, p-value_allelic_ = 1.82 × 10^−10^ and p-value_RA_ = 6.60 × 10^−9^. However, for this SNP 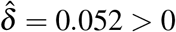 with p-value_HWE_ = 3.09 × 10^−4^; the HWE test is significant but not suggesting genotyping error. Thus, *T*_allelic_ is too optimistic. In contrast, for the third most significant SNP, 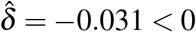 and p-value_HWE_ = 0.040. In that case, *T*_allelic_ is conservative while the proposed *T*_RA_ is not only robust but also more powerful, where p-value_allelic_ = 5.84 × 10^−6^ and p-value_RA_ = 8.86 × 10^−7^.

**Figure 3:**
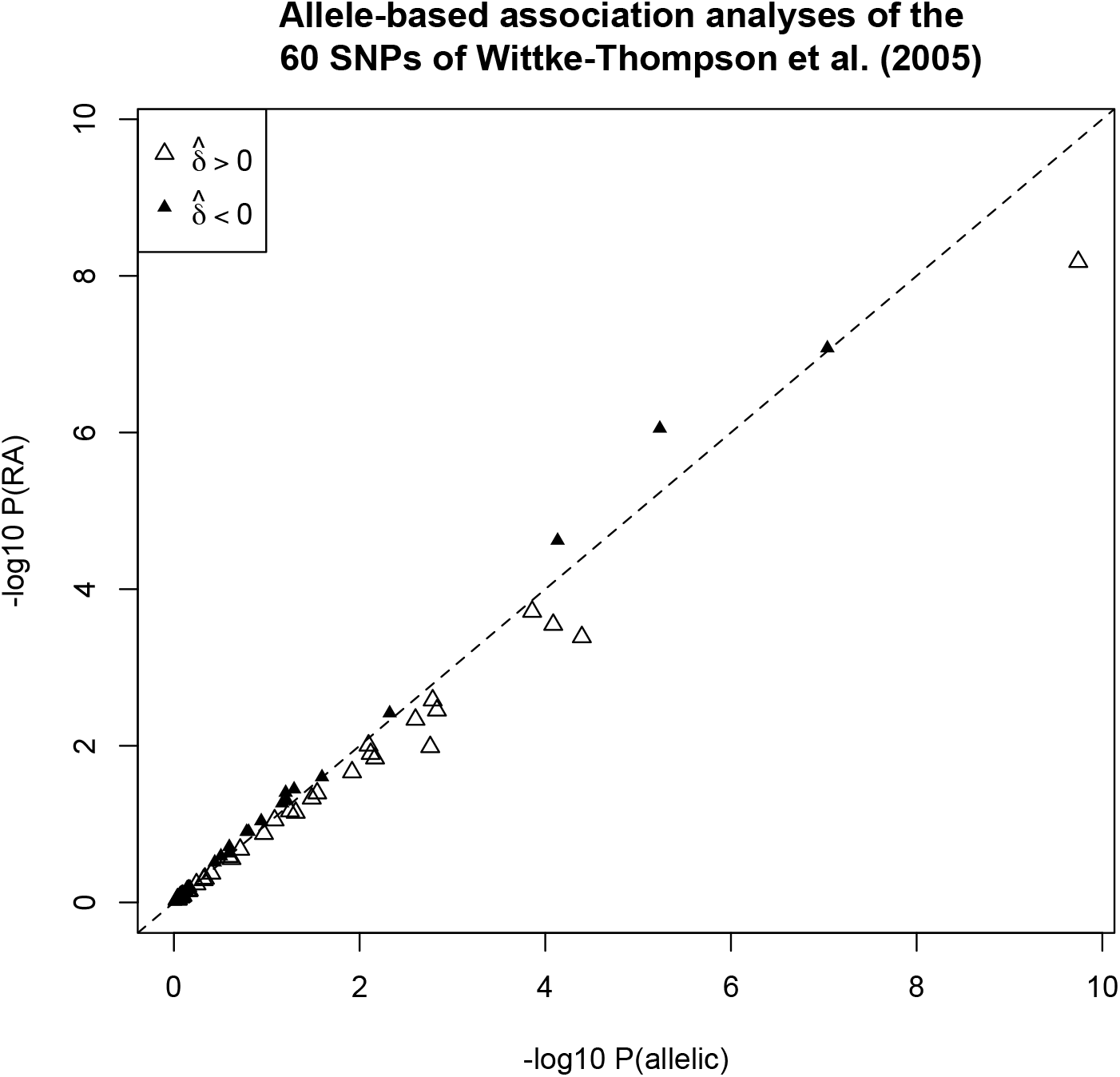
Results of application 1. Allele-based association tests of the 60 SNPs identified in Wittke-Thompson et al. (2005), contrasting the proposed RA method, *T*_RA_, with the classical allelic test, *T*_allelic_. Unfilled triangles are for SNPs with 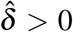 (*T*_allelic_ having inflated type 1 error), and filled triangles are for SNPs with 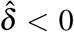 (*T*_allelic_ having deflated type 1 error); see Figure 1 for theoretical results and Figure 2 for simulation results regarding type 1 error control of the two methods.

### 5.3 Application 2 - a cystic fibrosis (CF) gene modifier study

Here we applied *T*_RA_ to simultaneously analyze two phenotypes using a sample with related individuals from the Canadian cystic fibrosis (CF) gene modifier study. The two phenotypes of interest are lung function, a quantitative trait (Taylor et al., 2011), and meconium ileus (MI), a binary trait (Gong et al., 2019). The sample of 2, 540 CF subjects includes 2, 420 singletons and 60 independent sib-pairs. For completeness, we first analyzed each phenotype individually using the proposed *allele-based* RA framework, and we compared the results with the traditional *genotype-based* method via the standard (generalized) linear mixed models (LMM or GLMM). We then analyzed both phenotypes simultaneously using *T*_RA_.

Figures 4(a) and 4(b) show that results of genotype-and allele-based methods are largely consistent; see Section 6 for a discussion. For the most significant SNP associated with MI (Figure 4(b)), the p-value of *T*_RA_ is 2.62 × 10^−6^, slightly smaller than 7.80 × 10^−6^ of the GLMM method. In addition, the proposed *T*_RA_ method can simultaneously analyzed both phenotypes and identify suggestive SNPs that have p-values several orders of magnitude smaller than those from studying one phenotype at a time (Figures 4(c) and 4(d)). However, these results are not genome-wide significant and establishing true association requires follow-up studies.

**Figure 4:**
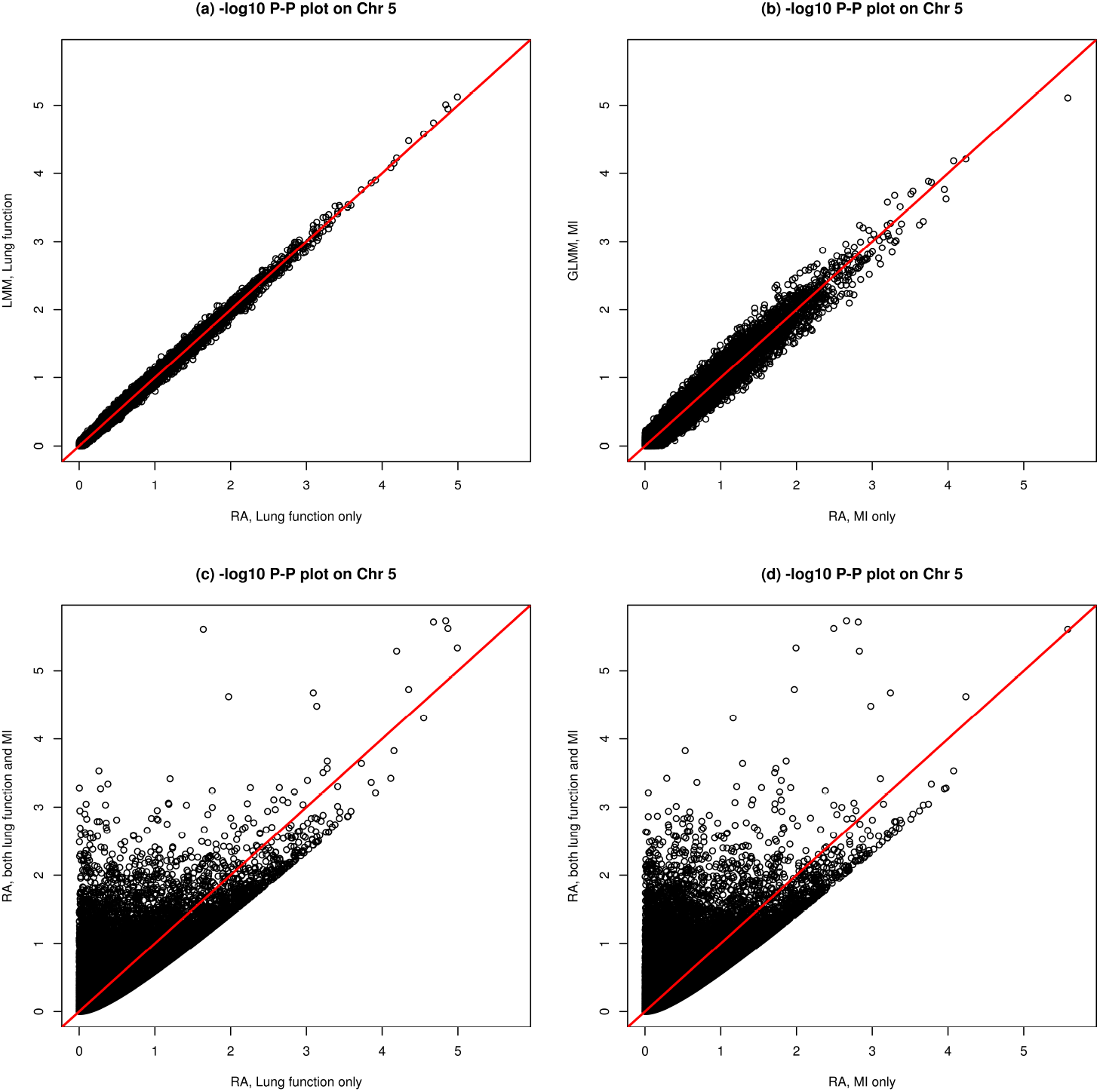
Results of application 2 - Chromosome 5-wide. Genetic association studies of lung function and meconium ileus of 34, 378 bi-allelic markers on chromosome 5, using a sample of 2, 540 individuals with cystic fibrosis of which 2, 420 are singletons and 120 are from 60 sib-pairs. LMM and GLMM are *genotype-based* association analyses based on, respectively, the standard linear mixed model for a continuous trait (i.e. lung) and generalized LMM for a binary trait (i.e. MI), and RA is the proposed allele-based association method that can also simultaneously analyze multiple traits using a sample of related individuals. Genome-wide results are shown in Web Figure 1.

Table 2 summarizes the association results for previously reported and replicated SNPs associated with CF lung function (Corvol et al., 2015) and MI association (Sun et al., 2012). For all the SNPs in Table 2, the proposed RA test yields slightly larger −log_10_(p-values) than LMM or GLMM, suggesting that the allele-based method has the potential to be more powerful than the traditional genotype-based approach. The RA simultaneous analysis of the two phenotypes, however, did not lead to more significant results; this is not surprising because these SNPs were selected based on the two previous single-phenotype analyses.

## 6 Discussion

To generalize the concept of comparing allele frequencies between case and control groups to more complex data settings, here we developed a novel robust allele-based (RA) regression framework that regresses individual alleles on phenotypes. Motivated by the earlier work of Chen (1983) for testing regression coefficients in independent samples, the proposed method utilizes the Gaussian model of (10) that (i) leads to a valid allelic association test through testing the regression coefficient *β*, (ii) adjusts for covariate effect through additional regression coefficient *γ*, (iii) explicitly models potential departure from Hardy–Weinberg equilibrium through *ρ* in the covariance matrix Σ, (iv) accounts for sample correlation through kinship coefficient *ϕ* in Σ, and (v) analyzes either a binary or a continuous phenotype, or multiple phenotypes simultaneously, where the phenotype data can be subjected to *Y* -dependent sampling.

The generalized allelic association test also unifies existing allelic tests. Under the HWE assumption and for a case-control study using an independent sample without covariates, the score test of *H*_0_ : *β* = 0 based on the simplified RA model (7) is identical to the classical allelic test in (2), *T*_RA, binary, *γ*=0*, ρ*=0_ = *T*_allelic_. In the presence of HWD, *T*_RA, binary, *γ*=0_ = *T*_allelic, Schaid_.

A crucial stage of this work is creating the two allele-based random variables, *G*_*i*1_ and *G*_*i*2_, and leveraging the ‘power’ of regression in new settings. The idea of reformulating an existing test statistic as a regression to facilitate method extension is not new. In their Reader Reaction to the generalized non-parametric Kurskal-Wallis test of Acar and Sun (2013) for handling group uncertainty, Wu and Guan (2015) presented “*a rank linear regression model and derived the proposed GKW statistic as a score test statistic*”. More recently, Soave and Sun (2017) showed that by first reformulating the original Levene’s test, testing for variance heterogeneity between *k* groups in an independent sample without group uncertainty, as a two-stage regression, the extension to more complex data is more straightforward.

The concept of ‘reverse’ regression has also been explored before, focusing on regressing *geno-type* on phenotype, notably by O’Reilly et al. (2012) for simultaneously analyses of multiple phenotypes. The corresponding MultiPhen method uses an ordinal logistic regression for the three genotype groups and then applies a likelihood ratio test in independent samples. Zhang et al. (2014) then examined GEE-based association tests for related individuals, but fundamentally these early tests are genotype-based.

Another stream of genotype-based ‘reverse’ or retrospective approach started with the quasi-likelihood method of Thornton and McPeek (2007) for case-control association testing with related individuals. The method first defines *X*_*i*_ = *G*_*i*_/2 ∈ {0, 1/2, 1}, then links the mean of *X*_*i*_ with *Y*_*i*_ via a logit transformation and uses the kinship coefficient matrix as the covariance matrix of *X*_*i*_, and finally obtains a quasi-likelihood score test. Subsequently, Feng (2014) and Feng et al. (2011) extended the method of Thornton and McPeek (2007) to a quasi-likelihood regression model that can incorporate multiple phenotypes. Although *E*(*X*_*i*_) = *E*(*G*_*i*_/2) can be interpreted as allele frequency, the quasi-likelihood score test is fundamentally a genotype-based association method. Further, the use of the kinship matrix alone as the covariance matrix requires the assumption of HWE.

Most existing family-based association studies rely on the *Y* -on-*G* prospective regression framework via LMM or GLMM (Eu-Ahsunthornwattana et al., 2014). For the application study in Section 5.3, we applied both the proposed RA method, and LMM (for the continuous CF lung function) and GLMM (for the binary meconium ileus status). Although there are differences in the (single-phenotype) analyses (Figures 4(a) and 4(b)), results are remarkably consistent. Interestingly, in the simplest case of an independent sample with no covariates, the corresponding RA test statistic has identical form as that derived from the genotype-based prospective regression model, as well as that from the non-parametric trend test (Web Appendix I). This inference similarity indirectly confirms the validity of the proposed approach (‘reverse’, allele-based and Gaussian model applied to a binary variable), but does not take away the contribution of this work. In particular, unlike LMM or GLMM, the proposed RA method can analyze multiple phenotype simultaneously as shown in Figures 4(c) and 4(d).

One of the challenges related to the proposed framework is the interpretation of parameter estimate for *β* even though its corresponding hypothesis testing is valid. Thus, we emphasize that the method developed here is tailored for rapid association testing not effect size estimation. We direct the readers to Krutchkoff (1967); Halperin (1970) for parameter estimation in the context of ‘reverse’ regression. The proposed RA framework provides a statistically efficient and computationally fast way for genome-wide association scans. For example, the analysis of Application 2 took 23.16 hours using one CPU core of an Intel Xeon Processor (Skylake, IBRS) at 2.40 GHz; the computation time is reduced to less than 2 hours when analyzing only the unrelated individuals in the CF sample.

Another difficulty present in any ‘reverse’ regression approach is the modeling and interpretation of gene-gene or gene-environment interactions. It is also not clear how to perform an allelic association test for an X-chromosomal variant; see Chen et al. (2018) for genotype-based association methods. Finally, although the proposed RA method has good type 1 error control when analyzing rare variants (Web Figure 2 and Web Table 2), power of the RA method in its current implementation, analyzing one SNP at a time, is likely to be low for a rare variant. Extending the RA methodology to simultaneously analyze multiple rare variants (Derkach et al., 2014) is of future interest but beyond the scope of this work.

The proposed framework, however, is flexible and promising in other directions. For example, the inclusion of parameter *ρ* in the RA model (7) is advantageous for both method comparison and further development. In the absence of *Y* and *Z* and sample correlation, we have shown that the score test of *H*_0_ : *ρ* = 0 is identical to the classical Pearson’s *χ*^2^ test of HWE in (1), *T*_HWE, RA_ = *T*_HWE, Pearson_. For more complex data, instead of developing individual remedies addressing specific challenges, the proposed method provides a principled approach for extensions. For example, we have shown in Section 4.1 that by introducing a population indicator we can derive a HWE test across populations that, in the simple setting of an independent sample, has identical form with that of Troendle and Yu (1994).

In terms of association testing, the value of explicitly modeling HWD via *ρ* in the regression is two fold. First, if there is a strong prior evidence for HWE, we can restrict *ρ* to be zero and establish a locally most powerful score test. Second, for the special case of a case-control study, Song and Elston (2006) and Wang and Shete (2010) have argued that departure from HWE in the case group provides additional evidence for association. However, the existing methods are ad-hoc. For example, the method of Song and Elston (2006) first conducts genotype-based association test and Pearson *χ*^2^ test of HWE separately, then aggregates the two (dependent) tests by a weighted sum, and finally evaluates the statistical significance via simulations. The proposed RA regression framework conceptually offers a new approach that could directly incorporate group-specific *ρ* into association inference, which we will explore as future work.

## Acknowledgment

The authors thank Dr. Lisa Strug and her lab for providing the cystic fibrosis application data. This research is funded by the Natural Sciences and Engineering Research Council of Canada (NSERC, RGPIN-04934 and RGPAS-522594), and the Canadian Institutes of Health Research (CIHR, MOP-310732) to LS. LZ is a trainee of the CANSSI-ONTARIO STAGE training program at the University of Toronto.

